# Decreased homotopic interhemispheric functional connectivity in children with autism spectrum disorder

**DOI:** 10.1101/2020.11.08.373126

**Authors:** Shuxia Yao, Menghan Zhou, Yuan Zhang, Feng Zhou, Qianqian Zhang, Zhongbo Zhao, Xi Jiang, Xiaolei Xu, Benjamin Becker, Keith M. Kendrick

**Author notes:** Corresponding authors **Correspondence:** Shuxia Yao or Keith M. Kendrick, No.2006, Xiyuan Ave., West Hi-Tech Zone, Chengdu, Sichuan 611731, China. or. **Lay Summary:** Homotopic interhemispheric functional connectivity plays an important role in synchronizing activity between the two hemispheres and is altered in adults and adolescents with autism spectrum disorder (ASD). In the present study focused on children with ASD, we have observed similar decreased homotopic connectivity in extensive brain networks, suggesting that alterations in homotopic interhemispheric connectivity may occur early in ASD and be a useful general biomarker across ages.

## Abstract

While a number of functional and structural changes occur in large-scale brain networks in autism spectrum disorder (ASD), reduced interhemispheric resting state functional connectivity (rsFC) between homotopic regions may be of particular importance as a biomarker. ASD is an early-onset developmental disorder and neural alterations are often age-dependent, reflecting dysregulated developmental trajectories, although no studies have investigated whether homotopic interhemispheric rsFC alterations occur in ASD children. The present study conducted a voxel-based homotopic interhemispheric rsFC analysis in 146 SD and 175 typically developing children under age 10 and examined associations with symptom severity in the Autism Brain Imaging Data Exchange datasets. Given the role of corpus callosum (CC) in interhemispheric connectivity and reported CC volume changes in ASD we additionally examined whether there were parallel volumetric changes in ASD children. Results demonstrated decreased homotopic rsFC in ASD children in the medial prefrontal cortex, precuneus and posterior cingulate cortex of the default mode network (DMN), the dorsal anterior cingulate cortex of the salience network, the precentral gyrus and inferior parietal lobule of the mirror neuron system, the lingual, fusiform and inferior occipital gyri of the visual processing network and thalamus. Symptom severity was associated with homotopic rsFC in regions in the DMN and visual processing network. There were no significant CC volume changes in ASD children. The present study shows that reduced homotopic interhemispheric rsFC in brain networks in ASD adults/adolescents is already present in children of 5-10 years old and further supports their potential use as a general ASD biomarker.

## Introduction

Autism spectrum disorder (ASD) is a heterogeneous neurodevelopmental disorder with core symptomatology generally characterized by deficits in social interaction, verbal and nonverbal communication and restrictive or stereotyped behavior (Kanner, 1943). Behavioral studies have revealed a variety of impairments in social-cognitive and affective domains in ASD patients, including deficits in joint attention (Charman, 2003), weak central coherence (Happé & Frith, 2006), avoidance of eye contact (Madipakkam, Rothkirch, Dziobek, & Sterzer, 2017), difficulties in face recognition (Fedor et al., 2018) and diminished interest and motivation in social interactions (Chevallier, Grèzes, Molesworth, Berthoz, & Happé, 2012; Morrison et al., 2017). Task-based neuroimaging studies have demonstrated that these deficits are mediated by dysfunctions in large-scale brain systems sub-serving social-cognitive and affective processes including the default mode network (DMN), mirror neuron system (MNS), theory of mind (ToM), and executive control networks (Dapretto et al., 2006; Herringshaw, Kumar, Rody, & Kana, 2018; Oberwelland et al., 2017; von dem Hagen, Stoyanova, Rowe, Baron-Cohen, & Calder, 2014; Wang, Dapretto, Hariri, Sigman, & Bookheimer, 2004).

The behavioral deficits in ASD and the need for diagnostics at an early age limit the application of task-based fMRI assessments in ASD. Consequently, resting state fMRI (rs-fMRI), which has the advantage of being task-free and providing comparable measures across different studies and sites, has been considered as a promising approach for investigating neural abnormalities and for providing possible promising biomarkers in ASD. In line with the task-based fMRI findings, ASD-associated alterations in the intrinsic organization of the brain as assessed by resting state functional connectivity (rsFC) have been found in the DMN, MNS, ToM and executive control networks (Chen, Yang, Wu, Chuang, & Huang, 2019; Fishman, Keown, Lincoln, Pineda, & Müller, 2014; Funakoshi et al., 2016; Monk et al., 2009), suggesting that large-scale aberrant functional organization occurs both in a task-free and task-based conditions in ASD. Additional brain networks showing rsFC abnormities in autistic patients have been reported in salience, mesolimbic reward and language networks (Lee, Park, James, Kim, & Park, 2017; Supekar et al., 2018; Uddin et al., 2013). Furthermore, atypical developmental trajectories in the functional and structural organization of these networks have also been found in ASD (Ecker, Bookheimer, & Murphy, 2015; Geschwind & Levitt, 2007; Washington et al., 2014).

Based on altered functional connectivity patterns, ASD has been proposed as a developmental dysfunctional connectivity syndrome with local overconnectivity but long-distance underconnectivity (Courchesne & Pierce, 2005; Geschwind & Levitt, 2007; Lau, Leung, & Lau, 2019). Recent studies have focused on interhemispheric functional connections particularly the homotopic ones, which may be of great importance in ASD. Homotopic interhemispheric functional connectivity is a measurement of the rsFC between each voxel in one hemisphere and its mirrored counterpart in the opposite hemisphere (Stark et al., 2008; Zuo et al., 2010). As a key characteristic of the intrinsic functional architecture of the brain, homotopic interhemispheric functional connections in the brain are exceptionally strong and may therefore play an important role in synchronizing activity between the two hemispheres (Stark et al., 2008). Anderson et al. (2011) first investigated the homotopic interhemispheric rsFC in ASD and found decreased homotopic interhemispheric rsFC between sensorimotor cortex, insula, fusiform gyrus, superior temporal gyrus, and superior parietal lobule in ASD compared to typically developing (TD) controls. Two other studies using the large Autism Brain Imaging Data Exchange (ABIDE) multisite datasets (ABIDE-I or -II), which have the advantage of greater statistical power, confirmed homotopic interhemispheric rsFC alterations in ASD, particularly in the interhemispheric communication of the posterior cingulate cortex (PCC), insula and thalamus (Di Martino et al., 2014 – ABIDE-I) and across a variety of large-scale networks (King et al., 2019; ABIDE-I and -II). Including gender as an additional covariate, Li et al. (2019) using the ABIDE-I dataset further confirmed decreased homotopic interhemispheric connectivity in ASD subjects and reported associations between interhemispheric functional connectivity in the PCC, insula and superior temporal gyrus and social communication impairment severity using the Autism Diagnostic Observation Schedule (ADOS). Thus, interhemispheric functional connectivity changes may represent a particular promising biomarker for discriminating ASD from TD individuals (Li et al., 2019). Note that some ASD-related rsFC changes are age-dependent (Hull, Jacokes, Torgerson, Irimia, & van Horn, 2017) and there is also evidence that homotopic interhemispheric connectivity shows different developmental trajectories in ASD compared with TD individuals (Kozhemiako et al., 2018). It is therefore possible that interhemispheric communication dysfunctions may also represent a result of dysregulated developmental trajectories. Although interhemispheric connectivity alterations have been investigated in ASD patients, subjects used in previous studies, including the ones using ABIDE datasets, are primarily high functioning adults/adolescents with ASD (e.g., mean = 22.4 years old in Anderson et al., 2011 and 17.39 years old for ABIDE-I). However, despite the neurodevelopmental nature of the disorder (Hull et al., 2017; Kozhemiako et al., 2018), no study to date has specifically examined homotopic interhemispheric rsFC changes in ASD children.

To shed new light on the utility of using homotopic interhemispheric rsFC alterations as biomarkers for ASD in general and as early biomarkers to determine ASD in childhood in particular, the present study has therefore conducted a voxel-based functional connectivity analysis to investigate whether alterations in homotopic interhemispheric rsFC are already present in ASD children and whether the changes are associated with ASD symptom severity. Furthermore, as the primary structure connecting the left and right brain hemispheres, corpus callosum (CC) volumes have been considered as an index of interhemispheric connectivity (Vidal et al., 2006; Wegiel et al., 2018) and volumetric CC alterations have also been associated with ASD (Anderson et al., 2011; Frazier & Hardan, 2009). The present study thus additionally examined whether there were parallel CC volume changes in ASD children and, if so, associations with symptom severity.

## Methods

### Participants

Resting-state fMRI and T1-weighted anatomical MRI images of 476 children under the age of 10 years (mean = 8.57, SD = 0.947; range: 5.3 - 10) were obtained from the ABIDE-I and -II datasets (http://fcon_1000.projects.nitrc.org/indi/abide/ for more information). The threshold of 10 years old was selected based on definition of age range for life phases from the American Academy of Pediatrics (Hagan, Shaw, & Duncan, 2008) and the World Health Organization (https://www.who.int/southeastasia/health-topics/adolescent-health). Consistent with Li et al. (2019), subjects were excluded in cases where preprocessing was not successful and head motion was excessive. Subjects were excluded in cases where: (a) fMRI images could not be successfully preprocessed, as determined by visual inspection of the normalization success; (b) mean framewise displacement (FD) was > 0.5 mm; (c) the percentage of FD > 0.5 mm was over 25%. After excluding subjects based on these criteria, data from sites with less than 4 subjects remaining were further deleted from the subsequent analyses. To further control for head motion, we matched head motion (mean FD) in the 2 groups (ASD group: mean = 0.20, SD = 0.096; control group: mean = 0.18, SD = 0.097; t(319) = 1.48, p = 0.139). These steps led to 146 subjects (50 from ABIDE-I) in the ASD group and 175 subjects (62 from ABIDE-I) in the TD control group being included in the final interhemispheric functional connectivity analysis, with the majority of subjects in the two groups (96 ASD vs. 113 control) being from the ABIDE-II dataset. Subject demographic information is reported in Supplementary Table S1 and clinical information of subjects with available scores on symptom severity as measured by ADOS (module 3) were reported in Supplementary Table S2 (N = 70). To increase transparency and replicability, a full list of subject IDs was also reported in Supplementary Table S3 and Table S4 for ABIDE-I and -II respectively. The data from the ABIDE datasets was fully anonymized. All contributions in this project were based on studies approved by local Institutional Review Boards. Written informed consent was obtained from all subjects or carers.

### Data Analyses

#### Interhemispheric rsFC Analyses

Interhemispheric rsFC analyses were similar to Li et al. (2019). Resting-state fMRI images were preprocessed using the Data Processing Assistant for Resting-State fMRI (DPARSF 2.3) software (Yan & Zang, 2010; http://www.restfmri.net) for each site separately following a standardized procedure (Yan et al., 2013). After discarding the first 10 images, pre-processing included slice timing correction, realignment, segmentation and spatial normalization. Segmentation was conducted based on a pediatric tissue probabilistic map created by the Template-O-Matic Toolbox while taking age and sex of children used in the present study into consideration (Wilke, Holland, Altaye, & Gaser, 2008). The images were then normalized to the Montreal Neurological Institute (MNI) template. Head-motion parameters (using the Friston 24-parameter model; Friston, Williams, Howard, Frackowiak, & Turner, 1996; Satterthwaite et al., 2013), global signal, white matter and cerebrospinal fluid were regressed out to control for potential confounding artifacts. To further exclude residual effects of head motion, we included the mean FD as a nuisance covariate in the group analysis (Power et al., 2014) and matched the mean FD between the 2 groups as described above. Following resampling with a 3 mm × 3 mm × 3 mm resolution, the pre-processed images were spatially smoothed with a Gaussian kernel (8-mm full-width at half-maximum) and filtered using a band pass filter (0.01-0.08 Hz). The homotopic interhemispheric functional connectivity map was then generated based on a voxel-mirrored homotopic connectivity (VMHC) analysis (Zuo et al., 2010) implemented in the DPARSF toolbox (Yan & Zang, 2010). As a voxel-based functional connectivity analysis approach, the VMHC reflects rsFC between each voxel in one hemisphere and its mirrored counterpart in the opposite hemisphere (Stark et al., 2008; Zuo et al., 2010). The homotopic interhemispheric functional connectivity maps were then converted into Z-maps using Fisher’s z transformation for subsequent group level analyses. To promote a strict control of center variability in the rsFC data the combatting batch effect (ComBat) harmonization method was applied, an approach that has been shown to remove unwanted variation associated with site/scanner while preserving inter-site biological variability (https://github.com/canlab/ComBatHarmonization#id-section2). ComBat models site-specific scaling factors and uses empirical Bayes to improve the estimation of site parameters for small sample sizes. Previous studies have demonstrated that the ComBat harmonization method is more advanced in controlling for site effects relative to conventional regression methods for different imaging modalities and that it performs well for multi-site imaging studies with only a few participants in each site or for unbalanced sample size between studies or sites (Fortin et al., 2017, 2018; Johnson, Li, & Rabinovic, 2007; Yu et al., 2018).

For the group level analyses, a two-sample t-test was used to test group functional connectivity differences between the ASD and TD children. To control for potential confounding effects, gender, age, total brain volume (grey matter + white matter) and head motion (mean FD) were included as nuisance covariates in the group analysis. A threshold of p < 0.05 false discovery rate (FDR) corrected for multiple comparisons at the voxel level was used in the whole-brain analysis and only clusters larger than 10 voxels were reported. To increase the sensitivity to determine whether the changes in homotopic interhemispheric rsFC observed in a sample predominatly including adolescents and adults (Li et al., 2019) already occur in ASD children, we further applied a mask encompassing regions that exhibited altered homotopic interhemispheric rsFC in these samples from Li et al. (2019). To this end a mask was created based on the thresholded statistic map (pFDR < 0.05; see Figure S1C) of group differences between ASD and TD control groups as reported in Li et al. (2019).

We also conducted correlation analyses between the homotopic interhemispheric connectivity strength and symptom severity using Pearson correlation coefficients. The homotopic interhemispheric connectivity strength was extracted from a 6-mm sphere centered on the peak voxel for regions showing altered homotopic interhemispheric connectivity in the ASD relative to the control group. ADOS consisted of social, communication, and stereotyped behavior subscales and total ADOS scores and scores for each subscale were available for 70 ASD children in the ABIDE datasets.

#### Corpus Callosum Volume Analyses

Volume analyses of the CC were conducted on T1 data using the VBM8 toolbox (http://dbm.neuro.unijena.de/vbm8/; Ashburner & Friston, 2000) and kept consistent with Li et al. (2019). Volume analyses of the CC were conducted on T1 data using the VBM8 toolbox (http://dbm.neuro.unijena.de/vbm8/; Ashburner and Friston, 2000) and kept consistent with Li et al. (2019). T1 images were normalized to the Montreal Neurological Institute (MNI) template and segmented into GM, WM and cerebrospinal fluid. Data quality check was performed via the modules of “display one slice for all images” and “check sample homogeneity using covariance”. 10 subjects were further excluded due to their overall covariance homogeneity being below 2 standard deviations from the mean. The remaining 139 ASD subjects and 172 controls were entered into the final analysis. Segmented WM images from these subjects were then smoothed with a Gaussian kernel of 8-mm full-width at half-maximum. Site effects on CC volumes were also controlled by employing the ComBat harmonization method (Fortin et al., 2017, 2018; Johnson, Li, & Rabinovic, 2007; Yu et al., 2018).

Group differences in CC volume were tested using a two-sample t-test on smoothed WM images. Gender, age and the total brain volume (TBV) were included as nuisance covariates to control for potential confounding effects. The structural CC mask was derived from the Talairach Daemon database atlas (Lancaster, 1997). Within the CC ROI, a threshold of p < 0.05 FDR corrected at peak-level was set for multiple comparisons and only clusters larger than 10 voxels reported.

## Results

### Interhemispheric Functional Connectivity

Two-sample t-tests were used to examine homotopic interhemispheric functional connectivity differences between the ASD and TD children. Analyses on the whole brain level revealed that ASD compared to TD children exhibited decreased homotopic interhemispheric functional connectivity in two clusters located in the PCC (MNI = 6, −45, 12, t = 4.88, pFDR < 0.05, voxels = 17; MNI = 6, −42, 24, t = 4.77, pFDR < 0.05, voxels = 25). Applying the mask generated from homotopic interhemispheric FC alterations previously observed in ASD adolescents and adults (Li et al., 2019) additionally revelaed decreased homotopic interhemispheric functional connectivity in regions of the DMN, including the PCC, the dorsal medial prefrontal cortex (mPFC), ventral mPFC, precuneus and the angular gyrus; the anterior cingulate cortex (ACC) of the salience network; the precentral gyrus and inferior parietal lobule (IPL) of the mirror neuron system; the visual processing network, including the lingual, fusiform, inferior and middle occipital gyri, inferior and middle temporal gyri; and thalamus (extending from PCC) (pFDR < 0.05; Figure 1 and Table 1) in ASD children. Categorization of regions into large-scale brain networks was based on previous reviews and studies on the default mode network (Raichle, 2015; Raichle et al., 2001; Seghier, 2013), the salience network (Menon & Uddin, 2010), the mirror neuron system (Iacoboni & Dapretto, 2006; Rizzolatti & Craighero, 2004), and the visual processing network (Bogousslavsky, Miklossy, Deruaz, Assal, & Regli, 2017; Eickenberg, Gramfort, Varoquaux, & Thirion, 2017; Machielsen, Rombouts, Barkhof, Scheltens, & Witter, 2000). Group difference on altered homotopic interhemispheric functional connectivity between ASD and TD control groups for each site was presented in supplemental Figure S2 using the PCC as an example. There were no regions showing significantly stronger interhemispheric functional connectivity in the ASD relative to TD groups using the same threshold (pFDR < 0.05).

**Table 1.** Regions showing decreased interhemispheric connectivity in the autism relative to the control groups (MNI coordinates).

| Brain Region | BA | No. Voxels | Peak t-value | x | y | z |
| --- | --- | --- | --- | --- | --- | --- |
| Posterior Cingulate Cortex | 31/7/23 | 753 | 4.88 | 6 | -45 | 12 |
| Posterior Cingulate Cortex |  |  | 4.77 | 6 | -42 | 24 |
| Precuneus |  |  | 3.91 | 6 | -60 | 27 |
| Lingual Gyrus |  |  | 3.84 | 27 | -54 | -6 |
| Midbrain |  |  | 3.47 | 3 | -27 | -6 |
| Fusiform |  |  | 3.16 | 30 | -69 | -3 |
| Thalamus |  |  | 2.93 | 12 | -30 | 3 |
| Middle Temporal Gyrus | 21 | 69 | 4.31 | 66 | -18 | -9 |
| Middle Occipital Gyrus | 18/19 | 106 | 4.28 | 30 | -99 | 3 |
| Inferior Occipital Gyrus |  |  | 3.10 | 33 | -90 | -6 |
| Anterior Cingulate Cortex | 24/32 | 41 | 4.05 | 6 | 36 | 3 |
| Ventral Medial Prefrontal Cortex |  |  | 2.27 | 6 | 36 | -9 |
| Inferior Temporal Gyrus | 37/20 | 32 | 4.02 | 66 | -54 | -12 |
| Middle Temporal Gyrus |  |  | 3.45 | 54 | -48 | -12 |
| Caudate |  | 63 | 3.61 | 15 | 6 | 18 |
| Dorsal Medial Prefrontal Cortex | 10 | 42 | 3.50 | 9 | 63 | 9 |
| Supramarginal/Angular Gyrus | 40 | 32 | 3.49 | 60 | -51 | 33 |
| Precentral Gyrus | 6 | 23 | 3.46 | 36 | -15 | 42 |
| Middle Frontal Gyrus | 8/6 | 84 | 3.46 | 27 | 18 | 51 |
| Postcentral Gyrus | 5 | 44 | 3.22 | 24 | -42 | 63 |
| Precuneus | 7 | 58 | 3.18 | 21 | -60 | 51 |
| Inferior Parietal Lobule |  |  | 2.50 | 36 | -60 | 48 |

All with a $p < 0.05$ FDR corrected threshold and voxels larger than 10 corrected within the mask from Li et al. (2019). BA: Brodmann Area. Brain coordinates are reported in the Montreal Neurological Institute (MNI) space.

**Figure 1.**
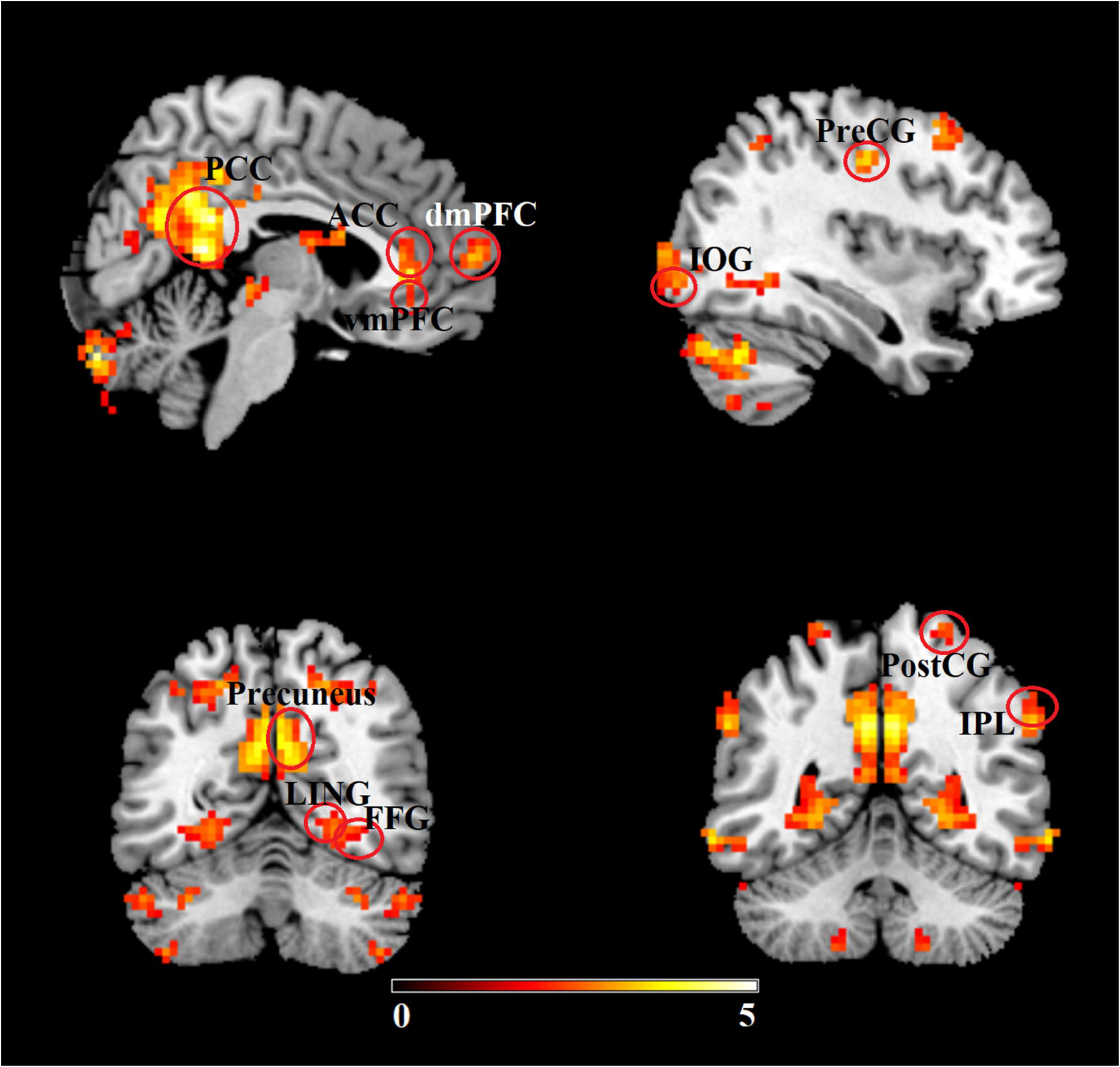
Regions showing significantly (pFDR < 0.05) decreased homotopic interhemispheric functional connectivity in the ASD compared to the typically developing groups. PCC = posterior cingulate cortex; dmPFC = dorsomedial prefrontal cortex; vmPFC = ventromedial prefrontal cortex; PreCG = precentral gyrus; IOG = inferior occipital gyrus; LING = lingual gyrus; FFG = fusiform gyrus; PostCG = postcentral gyrus; IPL = inferior parietal lobule. Color bar indicates t values of the statistical map.

These findings were consistent with the patterns found in our previous study (Li et al., 2019). Given that subjects from the ABIDE-I dataset used in the present study were a small part (~ 12.96%) of the subject sample used in our previous study (Li et al., 2019), we aimed at further controlling the possibility that subject overlap may have contributed to the consistent findings between the two studies. To this end, we further conducted an identical analysis using only ASD and TD children from the ABIDE-II dataset and found a highly similar but less robust pattern (see Figure S1A and B), which may be due to the smaller sample size when focusing only on the ABIDE-II dataset. Additionally, we also confirmed that removing data from children in the ABIDE-I dataset did not influence the pattern of results we had previously reported using data predominantly from adolescents and adults (Li et al., 2019) (see Figure S1C and D).

### Correlations between Interhemispheric Functional Connectivity and Symptom Severity

Correlation analyses between homotopic interhemispheric connectivity strength and symptom severity were first conducted for the total ADOS scores. Results revealed that the total ASD symptom load (total ADOS scores) showed a significant positive correlation with the homotopic interhemispheric functional connectivity strength of the PCC (MNI = 6, −45, 12; Pearson r = 0.268, p = 0.025; Figure 2A), but a negative correlation with the middle temporal gyrus (MTG; Pearson r = −0.240, p = 0.046; Figure 2B). Further explorations using sub-scale scores revealed significant positive correlations between the homotopic connectivity strengths of the PCC (MNI = 6, −45, 12) and social sub-scale scores (Pearson r = 0.254, p = 0.034; Figure 2C), and of the precuneus (Pearson r = 0.237, p = 0.048; Figure 2D) for the communication sub-scale scores. Homotopic functional connectivity strengths of the PCC (MNI = 6, −42, 24; Pearson r = 0.252, p = 0.035; Figure 2E), the inferior temporal gyrus (ITG; Pearson r = 0.274, p = 0.022; Figure 2F), the precuneus (Pearson r = 0.362, p = 0.002; Figure 2G) and the IPL (Pearson r = 0.242, p = 0.043; Figure 2H) were also positively associated with stereotyped behavior sub-scale scores. In contrast, significant negative correlations were found between homotopic connectivity strengths of the MTG (Pearson r = −0.250, p = 0.037; Figure 2I) and the inferior occipital gyrus (IOG; Pearson r = −0.237, p = 0.048; Figure 2J) and social sub-scale scores. However, none of these correlations can survive family-wise error correction for multiple comparisons (p < 0.05/22 = 0.002, where 22 is the total number of regions showing significant altered homotopic connectivity in ASD as presented in Table 1).

**Figure 2.**
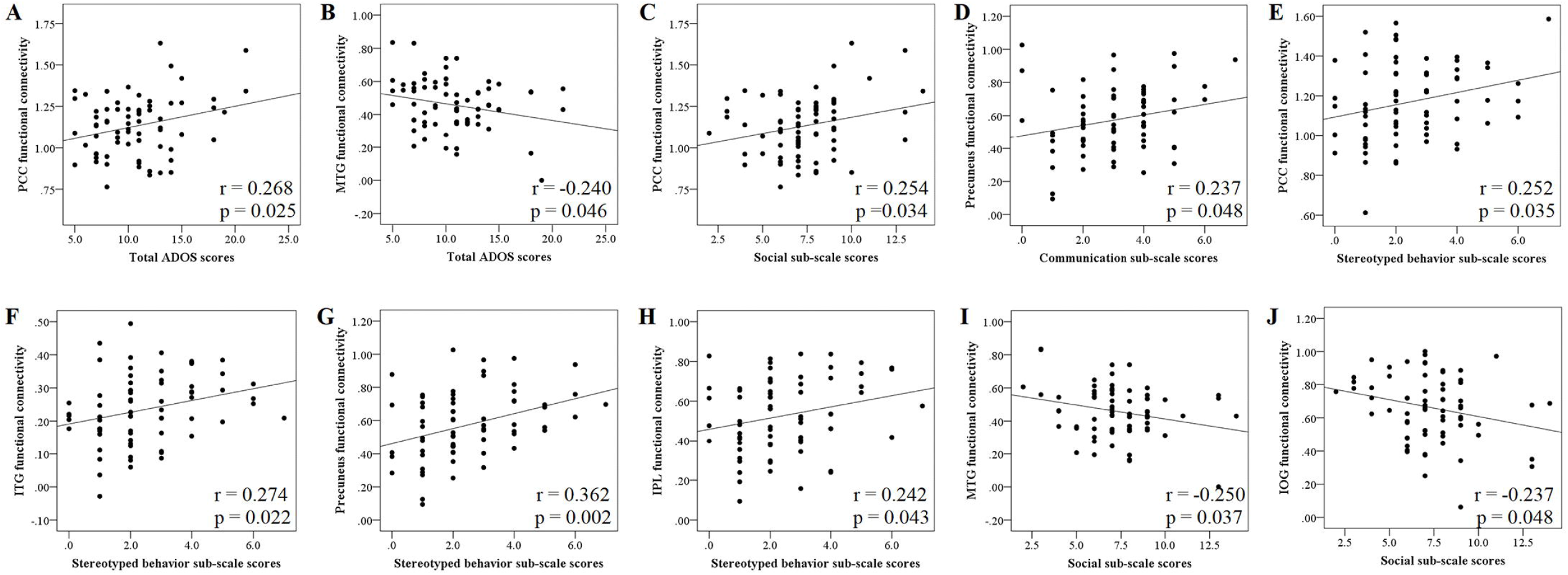
Correlations between the homotopic interhemispheric functional connectivity strengths of the posterior cingulate cortex (PCC) (A) and the middle temporal gyrus (MTG) (B) and ADOS total scores. Correlations between the homotopic connectivity strengths of the PCC and social sub-scale scores (C), and of the precuneus (D) for the communication sub-scale scores. A consistent positive correlation pattern was found between the homotopic connectivity of the PCC (E), the inferior temporal gyrus (F), the precuneus (G) and the inferior parietal lobule (H) and stereotyped behavior sub-scale scores. However, the homotopic connectivity strengths of the MTG (I) and inferior occipital gyrus (J) were negatively associated with social sub-scale scores.

### Corpus Callosum Volume

The ROI-based analysis of CC volume did not reveal any significant volume changes in the ASD compared to the TD group at a threshold of pFDR < 0.05.

## Discussion

The present study investigated homotopic interhemispheric rsFC alterations in ASD children under 10 years of age and their association with symptom severity. Given that the CC is the key fiber tract connecting the two hemispheres, changes in CC volumes were also examined. Results showed decreased homotopic interhemispheric functional connectivity in regions of the DMN, the salience network, the mirror neuron system, the visual processing network and thalamus in the ASD relative to the TD group, with interhemispheric functional connectivity of regions mainly in the DMN and the visual processing network being correlated with symptom severity. No significant CC volume changes were found in ASD children relative to TD controls. Altered homotopic functional connectivity may therefore be a potential biomarker for ASD in both children and adults.

Previous studies investigating homotopic interhemispheric rsFC changes in ASD mainly used subjects ranging from children to adults (Anderson et al., 2011; Di Martino et al., 2014; King et al., 2019; Li et al., 2019). For example, subjects from the ABIDE-I dataset used in most of these studies have an average age of 17.39 years with only 11.92% under the age of 10. However, only focusing on ASD children aged under 10, the present study using combined ABIDE-I and -II datasets found similar widespread decreased homotopic interhemispheric rsFC patterns as reported in these previous predominantly adult/adolescent-based studies. More specifically, the present study replicated the majority of brain networks exhibiting decreased homotopic interhemispheric connectivity in ASD patients reported by Li et al. (2019). These included the PCC, the dorsal and ventral mPFC, precuneus and the angular gyrus of the DMN, the dACC of the salience network, the precentral gyrus and IPL of the mirror neuron system, the lingual, fusiform and inferior occipital gyri of the visual processing network and thalamus. These networks are associated with a variety of cognitive and social/affective domains including self-referential/consciousness processing and theory of mind (the DMN; Davey, Pujol, & Harrison, 2016; Raichle, 2015; Saxe, Moran, Scholz, & Gabrieli, 2006), cognitive control and emotion regulation (dACC; MacDonald, Cohen, Stenger, & Carter, 2000; Stevens, Hurley, & Taber, 2011), action/intentions understanding and imitation (the mirror neuron system; Cattaneo & Rizzolatti, 2009; Oberman & Ramachandran, 2007) and the processing and integration of afferent sensory information (thalamus; Halassa & Kastner, 2017; Hwang, Bertolero, Liu, & D’Esposito, 2017). All of these systems can be dysfunctional in ASD (Burrows, Laird, & Uddin, 2016; Cheng, Rolls, Gu, Zhang, & Feng, 2015; Dichter, Felder, & Bodfish, 2009; Hamilton, 2013; Linke, Keehn, Pueschel, Fishman, & Müller, 2018; Padmanabhan, Lynch, Schaer, & Menon, 2017). The present study thus extends previous ones by providing the first evidence that, at least for children included in the ABIDE datasets, alterations in homotopic interhemispheric rsFC in large-scale brain networks have already occurred in individuals aged from 5 to 10 years of age.

Furthermore, correlation analyses between the homotopic interhemispheric functional connectivity and symptom severity revealed negative correlations between interhemispheric functional connectivity of the MTG and total ADOS scores and between homotopic interhemispheric connectivity of the MTG and the IOG and social sub-scale scores. This may indicate that reduced synchronization between the left and right parts of the MTG and the IOG, as reflected by lower interhemispheric functional connectivity, corresponds to greater symptom load in ASD. This is also consistent with negative correlation patterns as found in our previous study involving primarily adults/adolescents from the ABIDE-I dataset (Li et al., 2019) and may indicate a closer clinical relevance of the homotopic interhemispheric connectivity of these regions specifically in ASD children. In contrast, significant positive correlations were found between interhemispheric functional connectivity of the PCC and total ADOS and social sub-scale scores and between the precuneus and communication subscale scores. Similar positive correlation patterns were also observed for the homotopic interhemispheric connectivity of the PCC, the ITG, the precuneus and the IPL with the stereotyped behavior subscale scores, indicating that stronger synchronization between the left and right parts of these regions corresponds to greater symptom load in ASD. Thus, associations between homotopic functional connectivity and symptom severity appear to be more complex in ASD children, particularly the positive correlations with regions within the DMN, contrast to some extent with our previous study using primarily adult/adolescent subjects where we observed a negative association between homotopic interhemispheric PCC functional connectivity and the ADOS communication sub-scale (Li et al., 2019). A study on the developmental trajectories of homotopic functional connectivity strength in ASD children has reported that some DMN regions show an increase in connectivity during the first decade of life, followed subsequently by a maintained decrease during adolescence and adulthood (Kozhemiako et al., 2018). It is possible that initially a greater reduction in the homotopic functional connectivity of the PCC and precuneus may offer some compensatory benefit and reduced symptoms, but that subsequently in a more fully developed brain in adolescence/adulthood there is benefit in increasing the homotopic functional connectivity strength. This might explain the positive correlation with symptom severity during childhood but negative correlation during adolescence/adulthood and is in accord with positive correlations between homotopic interhemispheric connectivity of the MTG and the IOG in the visual processing network and ADOS scores as found in the present study and that homotopic functional connectivity strength of visual processing regions in ASD show flatter developmental trajectories relative to regions in DMN (Kozhemiako et al., 2018). Interestingly, the study on homotopic developmental trajectories also reported a positive association with ADOS scores as a function of developmental curvature coefficient (Kozhemiako et al., 2018). However, some caution is required when interpreting this finding since ADOS scores were only available for a relatively small number of ASD children in our current study.

Reduced homotopic functional connectivity has also been consistently reported in another developmental psychiatric disorder, schizophrenia (Guo et al., 2013; Li, Xu, Zhang, Hoptman, & Zuo, 2015) suggesting that impaired interhemispheric connectivity may be a hallmark of these disorders and contribute to some of the cognitive, social, motor and self-processing dysfunctions common to both. Decreased homotopic functional connectivity has been observed in early onset schizophrenia patients (mean age 14.5 years – Li et al., 2015) as well as adults (Guo et al., 2013), although specific regions associated with symptom severity differ from those associated with symptom severity in ASD.

We did not find significant differences between the ASD and TD children in CC volume. Previous findings on CC volume changes in ASD patients have been inconsistent, with some previous studies reporting reductions in both ASD adults and children (Haar, Berman, Behrmann, & Dinstein, 2016; Hardan et al., 2009; Keary et al., 2019; Li et al., 2019) and others in contrast observing no significant alterations (Herbert et al., 2014; Lefebvre, Beggiato, Bourgeron, & Toro, 2015; Rice et al., 2005). The inconsistency between these findings may well be as a result of many of the previous studies being statistical underpowered (Lefebvre et al., 2015) and CC volume changes derived from simple VBM-derived measures may also be too coarse an index to characterize subtle changes in ASD (Lefebvre et al., 2015; Li et al., 2019).

There are some limitations in the present study. Firstly, the ages of the ASD and TD children included in the present study ranged from 5 to 10 years old and therefore it is possible that younger children might not show similar reduced homotopic functional connectivity. This needs to be taken into consideration when making general inferences based on findings concerning age-independency of homotopic connectivity changes. Secondly, the ABIDE dataset only includes high-functioning ASD patients, therefore the current findings related to ASD children may not necessarily apply to low-functioning ASD populations. Finally, ADOS measurements were only available for a relatively small number of ASD children included in the study.

In conclusion, the present study using the ABIDE dataset has revealed decreased homotopic interhemispheric functional connectivity in diffuse brain networks including the DMN, the salience network, the mirror neuron system, the visual processing network, and thalamus in the ASD relative to the TD children between the ages of 5 and 10. Homo topic functional connectivity changes between regions in the DMN and the visual processing network were associated with the symptom severity. Thus, reduced homotopic interhemispheric functional organization appears to be present in both high functioning ASD children and adults/adolescents and may provide a useful general biomarker across both early and late developmental stages of the disorder.

## Acknowledgments

This study was supported by the National Natural Science Foundation of China (NSFC) grants (grant number: 31700998, 31530032) and by the Department of Science and Technology of Guangdong Province (grant number: 2018B030335001). The primary funding source for ABIDE-II is from NIMH 5R21MH107045.

