## Supplemental information for "Decreased homotopic interhemispheric functional connectivity in children with autism spectrum disorder"

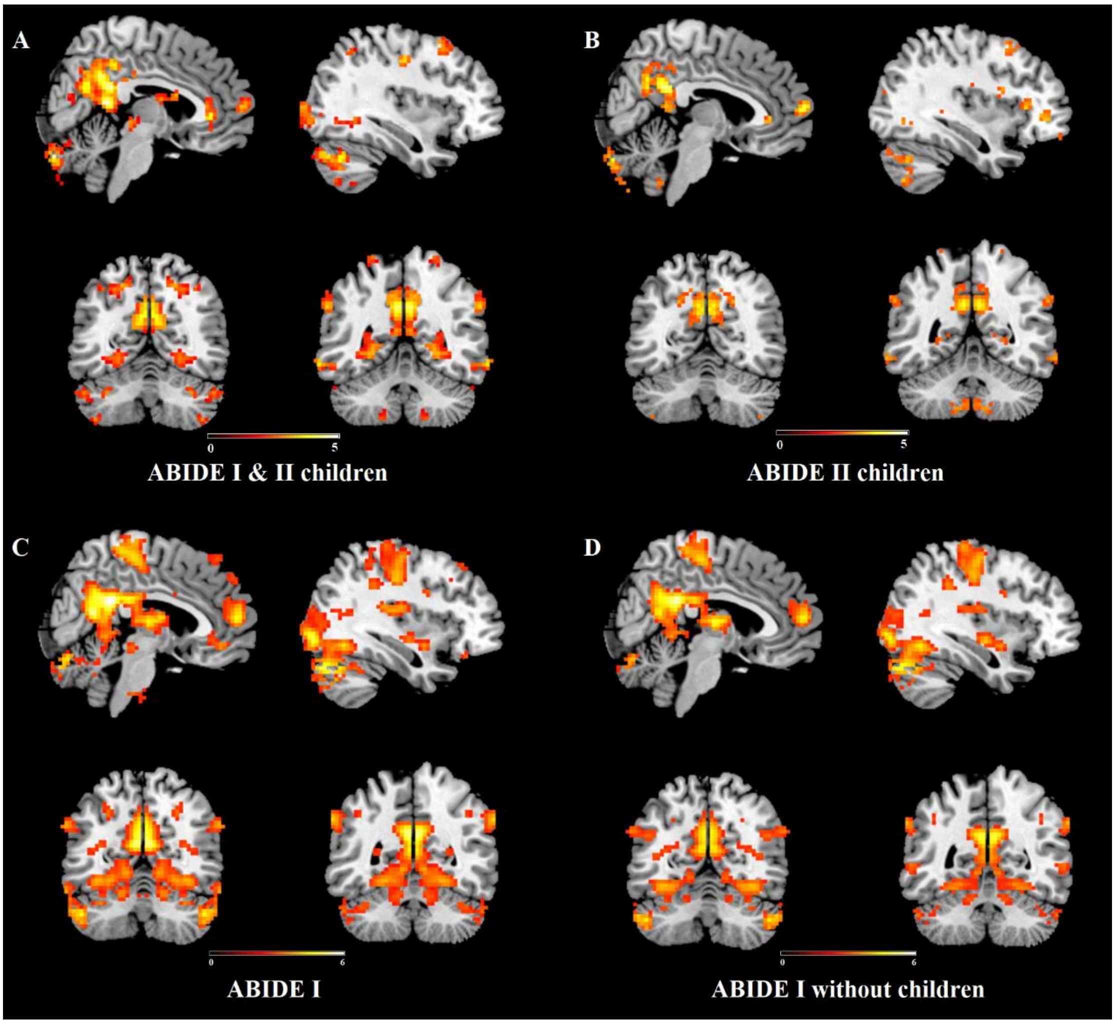


**Figure S1.** Regions showing altered homotopic interhemispheric functional connectivity using children subjects from both the ABIDE-I and -II datasets (A) and from only the ABIDE-II dataset (B). Regions showing altered homotopic interhemispheric functional connectivity using subjects from the ABIDE-I dataset found in Li et al. (2019) (C) and from the ABIDE-I dataset without ASD and TD children. All statistic maps were displayed with a $p < 0.05$ FDR corrected threshold except (B) where a $p < 0.01$ uncorrected threshold is used. Color bar indicates t values of the statistical map.

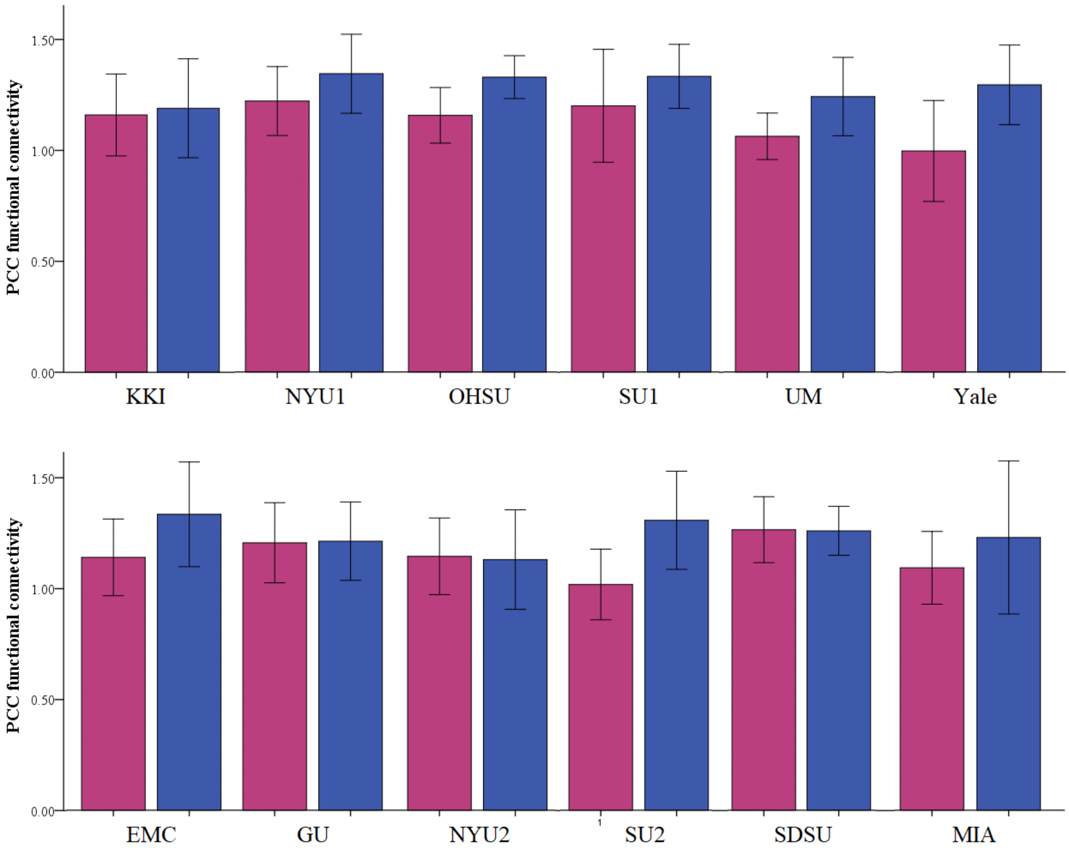


**Figure S2.** Group difference on altered homotopic interhemispheric functional connectivity between ASD and TD control groups for each site using the posterior cingulate cortex as an example. Given same scanner and scanning parameters were used, data from the KKI and OHSU were combined together. NYU 1, SU1, UM and Yale are from ABIDE-I dataset and EMC, GU, NYU2, SU2, SDSU and MIA are from ABIDE-II dataset.

**Table S1.** The demographic of subjects satisfying inclusion criteria in the functional connectivity analysis.

| ABIDE-I | Autism | | | | | | Control | | | | | |
|---|---|---|---|---|---|---|---|---|---|---|---|---|
| | Subjects | Age | M/F | FIQ | VIQ | PIQ | Subjects | Age | M/F | FIQ | VIQ | PIQ |
| KKI | 6 | 8.30±0.16 | 4/2 | 96.83±18.27 | / | / | 17 | 9.18±0.61 | 11/6 | 113.29±7.92 | / | / |
| NYU | 21 | 8.74±0.99 | 21/0 | 112.00±20.25 | 105.48±16.04 | 116.95±22.61 | 17 | 8.30±0.86 | 11/6 | 115.29±12.59 | 115.94±14.85 | 111.29±11.56 |
| OHSU | 4 | 9.22±0.82 | 4/0 | 103.73±22.52 | / | / | 5 | 9.23±0.62 | 5/0 | 116.08±10.69 | / | / |
| SU | 10 | 8.83±0.76 | 8/2 | 119.80±15.57 | 115.30±19.76 | 120.10±16.46 | 12 | 8.81±0.61 | 9/3 | 110.67±18.40 | 108.50±22.83 | 110.67±18.82 |
| UM | 4 | 9.58±0.25 | 3/1 | 99.38±32.45 | 112±45.89 | 86.75±19.02 | 5 | 9.40±0.68 | 3/2 | 100.90±5.31 | 109.20±7.53 | 92.60±8.65 |
| Yale | 5 | 8.52±1.32 | 4/1 | 94.80±12.07 | 97.80±13.29 | 93±12.65 | 6 | 8.90±0.81 | 5/1 | 117.17±14.78 | 110.50±13.37 | 114.67±14.25 |
| **ABIDE-II** | **Autism** | | | | | | **Control** | | | | | |
| | **Subjects** | **Age** | **M/F** | **FIQ** | **VIQ** | **PIQ** | **Subjects** | **Age** | **M/F** | **FIQ** | **VIQ** | **PIQ** |
| EMC | 14 | 8.32±0.94 | 11/3 | / | / | 102.18±12.16 | 16 | 7.99±0.83 | 13/3 | / | / | 99.07±15.23 |
| GU | 9 | 9.28±0.69 | 7/2 | 123.25±14.28 | 128.50±13.93 | 121.20±17.63 | 19 | 8.77±0.43 | 13/6 | 122.26±16.70 | 121.37±17.79 | 118.42±14.80 |
| KKI | 17 | 8.99±0.55 | 10/7 | 97.44±14.30 | 106.24±18.24 | 101.12±13.87 | 26 | 8.78±0.42 | 16/10 | 112.08±10.04 | 116.73±13.21 | 110.15±13.71 |
| NYU | 34 | 7.47±1.05 | 32/2 | 103.03±18.28 | 104.97±17.38 | 104.32±21.77 | 19 | 7.89±1.29 | 18/1 | 113.00±14.60 | 116.00±16.30 | 108.00±14.67 |
| OHSU | 7 | 8.14±0.90 | 5/2 | 102.57±23.15 | / | / | 16 | 8.56±0.51 | 10/6 | 119.50±13.48 | / | / |
| SDSU | 5 | 9.12±0.68 | 4/1 | 107.80±15.06 | 103.00±18.56 | 111.60±20.72 | 4 | 9.10±0.85 | 4/0 | 113.75±9.54 | 109.25±12.74 | 115.00±6.53 |
| SU | 4 | 9.53±0.74 | 4/0 | 118.75±21.09 | 122.75±13.82 | 111.00±23.02 | 6 | 9.34±0.47 | 5/1 | 112.00±12.13 | 117.33±20.50 | 104.67±9.93 |
| MIA | 6 | 8.44±0.91 | 4/2 | 97.83±15.83 | 95.83±15.89 | 103.33±15.91 | 7 | 8.19±0.86 | 5/2 | 109.86±14.08 | 108.29±7.46 | 107.14±23.14 |
| **Mean/total** | 146 | 8.48±1.07 | 121/25 | 106.03±19.40 | 107.80±19.37 | 107.27±20.15 | 175 | 8.62±0.84 | 128/47 | 114.43±13.40 | 115.09±15.89 | 109.35±15.40 |

FIQ: Full IQ; VIQ: Verbal IQ; PIQ: Performance IQ. Mean/total for subjects indicates the total number of subjects; others indicate mean ± SD. FIQ scores are available for 130 autistic children and 159 controls. VIQ scores are available for 114 autistic children and 121 controls. PIQ scores are available for 122 autistic children and 135 controls. Given that same scanning parameters were used for OHSU in ABIDE I and II, data of OHSU was treated as one site.

**Table S2.** The clinical information of subjects in the brain-autistic symptoms correlation analysis.

| | Subjects | ADOS TOTAL | ADOS COMM | ADOS SOCIAL | STEREO BEHAV |
|---|---|---|---|---|---|
| **ABIDE-I** | | | | | |
| KKI | 6 | 11.00±2.37 | 3.50±1.38 | 7.50±1.22 | 3.50±1.05 |
| NYU | 21 | 10.86±4.35 | 3.29±1.82 | 7.57±3.04 | 2.95±1.77 |
| OHSU | 4 | 10.00±3.16 | 3.50±1.29 | 6.50±2.08 | 1.75±0.95 |
| SU | / | / | / | / | / |
| UM | | | | | |
| Yale | / | / | / | / | / |
| **ABIDE-II** | | | | | |
| EMC | / | / | / | / | / |
| GU | 7 | 9.71±3.30 | 2.86±1.07 | 6.86±2.97 | 1.43±1.40 |
| KKI | 10 | 14.80±3.29 | 3.40±1.26 | 8.40±2.01 | 3.00±1.70 |
| NYU | 15 | 8.73±2.52 | 2.27±1.39 | 6.47±1.51 | 1.93±1.39 |
| OHSU | 3 | 10.33±2.08 | 3.33±1.15 | 7.00±1.00 | 2.33±1.53 |
| SDSU | / | / | / | / | / |
| SU | 4 | 10.75±2.87 | 2.25±1.26 | 8.50±1.73 | 2.00±1.41 |
| MIA | / | / | / | / | / |
| **Mean/total** | 70 | 10.79±3.73 | 3.01±1.49 | 7.34±2.32 | 2.49±1.59 |

**ADOS TOTAL:** Classic Total Autism Diagnostic Observation Schedule (ADOS) Score (Communication Subscore + Social Interaction Subscore); **ADOS_COMM:** Communication Total Subscore of the Classic ADOS; **ADOS SOCIAL:** Social Total Subscore of the Classic ADOS; **ADOS STEREO BEHAV:** Stereotyped Behaviors Total Subscore of the Classic ADOS; **ADOS_ALL:** Total Score of All the Three Subscale Including Communication, Social, and Stereotyped Behaviors Subscales. Mean/total for subjects indicates the total number of subjects.

**Table S3.** IDs of subject from ABIDE-I.

| Autism group | | Control group | |
|---|---|---|---|
| **SITE_ID** | **SUB_ID** | **SITE_ID** | **SUB_ID** |
| KKI | 50792 | KKI | 50776 |
| KKI | 50795 | KKI | 50777 |
| KKI | 50802 | KKI | 50778 |
| KKI | 50803 | KKI | 50779 |
| KKI | 50804 | KKI | 50780 |
| KKI | 50825 | KKI | 50781 |
| NYU | 50965 | KKI | 50784 |
| NYU | 50967 | KKI | 50786 |
| NYU | 50968 | KKI | 50789 |
| NYU | 50969 | KKI | 50790 |
| NYU | 50970 | KKI | 50809 |
| NYU | 50977 | KKI | 50812 |
| NYU | 50978 | KKI | 50814 |
| NYU | 50979 | KKI | 50816 |
| NYU | 50980 | KKI | 50817 |
| NYU | 50982 | KKI | 50819 |
| NYU | 50986 | KKI | 50820 |
| NYU | 50987 | NYU | 51036 |
| NYU | 50989 | NYU | 51038 |
| NYU | 51002 | NYU | 51039 |
| NYU | 51003 | NYU | 51040 |
| NYU | 51010 | NYU | 51041 |
| NYU | 51011 | NYU | 51042 |
| NYU | 51012 | NYU | 51064 |
| NYU | 51013 | NYU | 51069 |
| NYU | 51032 | NYU | 51070 |
| NYU | 51033 | NYU | 51078 |
| OHSU | 50146 | NYU | 51079 |
| OHSU | 50150 | NYU | 51080 |
| OHSU | 50152 | NYU | 51081 |
| OHSU | 50153 | NYU | 51082 |
| STANFORD | 51160 | NYU | 51083 |
| STANFORD | 51162 | NYU | 51084 |
| STANFORD | 51163 | NYU | 51085 |
| STANFORD | 51164 | OHSU | 50161 |
| STANFORD | 51169 | OHSU | 50163 |
| STANFORD | 51172 | OHSU | 50165 |
| STANFORD | 51173 | OHSU | 50166 |
| STANFORD | 51174 | OHSU | 50168 |
| STANFORD | 51175 | STANFORD | 51180 |

| | | | |
|---|---|---|---|
| STANFORD | 51176 | STANFORD | 51181 |
| UM_1 | 50300 | STANFORD | 51183 |
| UM_1 | 50307 | STANFORD | 51184 |
| UM_1 | 50310 | STANFORD | 51185 |
| UM_1 | 50318 | STANFORD | 51186 |
| YALE | 50609 | STANFORD | 51187 |
| YALE | 50617 | STANFORD | 51188 |
| YALE | 50622 | STANFORD | 51189 |
| YALE | 50625 | STANFORD | 51190 |
| YALE | 50627 | STANFORD | 51191 |
| | | STANFORD | 51193 |
| | | UM_1 | 50357 |
| | | UM_1 | 50358 |
| | | UM_1 | 50363 |
| | | UM_1 | 50366 |
| | | UM_1 | 50374 |
| | | YALE | 50553 |
| | | YALE | 50554 |
| | | YALE | 50560 |
| | | YALE | 50564 |
| | | YALE | 50566 |
| | | YALE | 50576 |

**Table S4.** IDs of subjects from ABIDE-II.

| Autism group | | Control group | |
|---|---|---|---|
| **SITE_ID** | **SUB_ID** | **SITE_ID** | **SUB_ID** |
| ABIDEII-EMC_1 | 29864 | ABIDEII-EMC_1 | 29891 |
| ABIDEII-EMC_1 | 29866 | ABIDEII-EMC_1 | 29892 |
| ABIDEII-EMC_1 | 29867 | ABIDEII-EMC_1 | 29896 |
| ABIDEII-EMC_1 | 29869 | ABIDEII-EMC_1 | 29899 |
| ABIDEII-EMC_1 | 29870 | ABIDEII-EMC_1 | 29901 |
| ABIDEII-EMC_1 | 29872 | ABIDEII-EMC_1 | 29904 |
| ABIDEII-EMC_1 | 29874 | ABIDEII-EMC_1 | 29905 |
| ABIDEII-EMC_1 | 29875 | ABIDEII-EMC_1 | 29906 |
| ABIDEII-EMC_1 | 29876 | ABIDEII-EMC_1 | 29907 |
| ABIDEII-EMC_1 | 29877 | ABIDEII-EMC_1 | 29908 |
| ABIDEII-EMC_1 | 29879 | ABIDEII-EMC_1 | 29909 |
| ABIDEII-EMC_1 | 29881 | ABIDEII-EMC_1 | 29911 |
| ABIDEII-EMC_1 | 29882 | ABIDEII-EMC_1 | 29912 |
| ABIDEII-EMC_1 | 29885 | ABIDEII-EMC_1 | 29913 |
| ABIDEII-GU_1 | 28754 | ABIDEII-EMC_1 | 29915 |
| ABIDEII-GU_1 | 28758 | ABIDEII-EMC_1 | 29916 |
| ABIDEII-GU_1 | 28778 | ABIDEII-GU_1 | 28746 |
| ABIDEII-GU_1 | 28790 | ABIDEII-GU_1 | 28759 |
| ABIDEII-GU_1 | 28805 | ABIDEII-GU_1 | 28763 |
| ABIDEII-GU_1 | 28811 | ABIDEII-GU_1 | 28767 |
| ABIDEII-GU_1 | 28815 | ABIDEII-GU_1 | 28770 |
| ABIDEII-GU_1 | 28822 | ABIDEII-GU_1 | 28776 |
| ABIDEII-GU_1 | 28825 | ABIDEII-GU_1 | 28783 |
| ABIDEII-KKI_1 | 29273 | ABIDEII-GU_1 | 28793 |
| ABIDEII-KKI_1 | 29274 | ABIDEII-GU_1 | 28801 |
| ABIDEII-KKI_1 | 29275 | ABIDEII-GU_1 | 28803 |
| ABIDEII-KKI_1 | 29276 | ABIDEII-GU_1 | 28804 |
| ABIDEII-KKI_1 | 29278 | ABIDEII-GU_1 | 28814 |
| ABIDEII-KKI_1 | 29281 | ABIDEII-GU_1 | 28827 |
| ABIDEII-KKI_1 | 29283 | ABIDEII-GU_1 | 28829 |
| ABIDEII-KKI_1 | 29285 | ABIDEII-GU_1 | 28836 |
| ABIDEII-KKI_1 | 29287 | ABIDEII-GU_1 | 28841 |
| ABIDEII-KKI_1 | 29293 | ABIDEII-GU_1 | 28842 |
| ABIDEII-KKI_1 | 29392 | ABIDEII-GU_1 | 28845 |
| ABIDEII-KKI_1 | 29404 | ABIDEII-IP_1 | 29596 |
| ABIDEII-KKI_1 | 29415 | ABIDEII-KKI_1 | 29295 |
| ABIDEII-KKI_1 | 29416 | ABIDEII-KKI_1 | 29307 |
| ABIDEII-KKI_1 | 29434 | ABIDEII-KKI_1 | 29309 |
| ABIDEII-KKI_1 | 29479 | ABIDEII-KKI_1 | 29310 |
| ABIDEII-KKI_1 | 29481 | ABIDEII-KKI_1 | 29313 |

| | | | |
|---|---|---|---|
| ABIDEII-NYU_1 | 29179 | ABIDEII-KKI_1 | 29318 |
| ABIDEII-NYU_1 | 29181 | ABIDEII-KKI_1 | 29324 |
| ABIDEII-NYU_1 | 29183 | ABIDEII-KKI_1 | 29332 |
| ABIDEII-NYU_1 | 29184 | ABIDEII-KKI_1 | 29334 |
| ABIDEII-NYU_1 | 29185 | ABIDEII-KKI_1 | 29339 |
| ABIDEII-NYU_1 | 29186 | ABIDEII-KKI_1 | 29340 |
| ABIDEII-NYU_1 | 29188 | ABIDEII-KKI_1 | 29343 |
| ABIDEII-NYU_1 | 29189 | ABIDEII-KKI_1 | 29364 |
| ABIDEII-NYU_1 | 29191 | ABIDEII-KKI_1 | 29386 |
| ABIDEII-NYU_1 | 29193 | ABIDEII-KKI_1 | 29387 |
| ABIDEII-NYU_1 | 29194 | ABIDEII-KKI_1 | 29390 |
| ABIDEII-NYU_1 | 29195 | ABIDEII-KKI_1 | 29395 |
| ABIDEII-NYU_1 | 29199 | ABIDEII-KKI_1 | 29402 |
| ABIDEII-NYU_1 | 29201 | ABIDEII-KKI_1 | 29407 |
| ABIDEII-NYU_1 | 29202 | ABIDEII-KKI_1 | 29418 |
| ABIDEII-NYU_1 | 29206 | ABIDEII-KKI_1 | 29424 |
| ABIDEII-NYU_1 | 29208 | ABIDEII-KKI_1 | 29439 |
| ABIDEII-NYU_1 | 29209 | ABIDEII-KKI_1 | 29446 |
| ABIDEII-NYU_1 | 29210 | ABIDEII-KKI_1 | 29460 |
| ABIDEII-NYU_1 | 29213 | ABIDEII-KKI_1 | 29469 |
| ABIDEII-NYU_1 | 29220 | ABIDEII-KKI_1 | 29470 |
| ABIDEII-NYU_2 | 29152 | ABIDEII-NYU_1 | 29225 |
| ABIDEII-NYU_2 | 29153 | ABIDEII-NYU_1 | 29226 |
| ABIDEII-NYU_2 | 29154 | ABIDEII-NYU_1 | 29227 |
| ABIDEII-NYU_2 | 29155 | ABIDEII-NYU_1 | 29228 |
| ABIDEII-NYU_2 | 29157 | ABIDEII-NYU_1 | 29229 |
| ABIDEII-NYU_2 | 29159 | ABIDEII-NYU_1 | 29230 |
| ABIDEII-NYU_2 | 29162 | ABIDEII-NYU_1 | 29231 |
| ABIDEII-NYU_2 | 29163 | ABIDEII-NYU_1 | 29232 |
| ABIDEII-NYU_2 | 29165 | ABIDEII-NYU_1 | 29235 |
| ABIDEII-NYU_2 | 29168 | ABIDEII-NYU_1 | 29236 |
| ABIDEII-NYU_2 | 29169 | ABIDEII-NYU_1 | 29237 |
| ABIDEII-NYU_2 | 29173 | ABIDEII-NYU_1 | 29238 |
| ABIDEII-NYU_2 | 29175 | ABIDEII-NYU_1 | 29241 |
| ABIDEII-OHSU_1 | 28925 | ABIDEII-NYU_1 | 29244 |
| ABIDEII-OHSU_1 | 28934 | ABIDEII-NYU_1 | 29245 |
| ABIDEII-OHSU_1 | 28936 | ABIDEII-NYU_1 | 29247 |
| ABIDEII-OHSU_1 | 28940 | ABIDEII-NYU_1 | 29251 |
| ABIDEII-OHSU_1 | 28968 | ABIDEII-NYU_1 | 29253 |
| ABIDEII-OHSU_1 | 28995 | ABIDEII-NYU_1 | 29254 |
| ABIDEII-OHSU_1 | 28998 | ABIDEII-OHSU_1 | 28956 |
| ABIDEII-SDSU_1 | 28869 | ABIDEII-OHSU_1 | 28961 |
| ABIDEII-SDSU_1 | 28875 | ABIDEII-OHSU_1 | 28969 |

| | | | |
|---|---|---|---|
| ABIDEII-SDSU_1 | 28879 | ABIDEII-OHSU_1 | 28970 |
| ABIDEII-SDSU_1 | 28901 | ABIDEII-OHSU_1 | 28973 |
| ABIDEII-SDSU_1 | 28905 | ABIDEII-OHSU_1 | 28977 |
| ABIDEII-SU_2 | 30171 | ABIDEII-OHSU_1 | 28978 |
| ABIDEII-SU_2 | 30173 | ABIDEII-OHSU_1 | 28979 |
| ABIDEII-SU_2 | 30179 | ABIDEII-OHSU_1 | 28980 |
| ABIDEII-SU_2 | 30186 | ABIDEII-OHSU_1 | 28983 |
| ABIDEII-U_MIA_1 | 29739 | ABIDEII-OHSU_1 | 28984 |
| ABIDEII-U_MIA_1 | 29740 | ABIDEII-OHSU_1 | 28990 |
| ABIDEII-U_MIA_1 | 30229 | ABIDEII-OHSU_1 | 29001 |
| ABIDEII-U_MIA_1 | 30238 | ABIDEII-OHSU_1 | 30155 |
| ABIDEII-U_MIA_1 | 30239 | ABIDEII-OHSU_1 | 30158 |
| ABIDEII-U_MIA_1 | 30243 | ABIDEII-OHSU_1 | 30163 |
| | 30248 | ABIDEII-SDSU_1 | 28852 |
| | 30254 | ABIDEII-SDSU_1 | 28863 |
| | | ABIDEII-SDSU_1 | 28868 |
| | | ABIDEII-SDSU_1 | 28881 |
| | | ABIDEII-SU_2 | 30194 |
| | | ABIDEII-SU_2 | 30197 |
| | | ABIDEII-SU_2 | 30198 |
| | | ABIDEII-SU_2 | 30205 |
| | | ABIDEII-SU_2 | 30207 |
| | | ABIDEII-SU_2 | 30208 |
| | | ABIDEII-U_MIA_1 | 30236 |
| | | ABIDEII-U_MIA_1 | 30242 |
| | | ABIDEII-U_MIA_1 | 30244 |
| | | ABIDEII-U_MIA_1 | 30246 |
| | | ABIDEII-U_MIA_1 | 30253 |
| | | ABIDEII-U_MIA_1 | 30255 |
| | | ABIDEII-U_MIA_1 | 30256 |
